# Effectiveness of Outer Hair Cells as Cochlear Amplifier: Coupled Oscillator Models

**DOI:** 10.1101/2025.08.02.668240

**Authors:** Kuni H. Iwasa

**Affiliations:** NIDCD, National Institutes of Health Bethesda, MD 20892, USA

**Keywords:** hearing, sensitivity, coupling oscillators, mechanics

## Abstract

Outer hair cells (OHCs) are essential for the sensitivity and frequency specificity of the mammalian ear. To perform this function, OHCs need to amplify the motion of the basilar membrane, which is much stiffer than themselves. OHCs must overcome this impedance mismatch for their amplifying function particularly at high frequencies, where the mismatch is largest. This issue could be solved by the existence of multiple modes of motion. Here, systems of two coupled oscillators are examined as the simplest of such cases. It is found that some of these model systems indeed make OHCs function as an effective amplifier by overcoming the impedance mismatch. This result suggests that the elaborate structure of the organ of Corti, which can support multiple modes of motion, is a key to the high frequency performance of the mammalian ear.

**Significance:** The mammalian ear depends on outer hair cells, which work as the cochlear amplifier. The mechanism, with which outer hair cells perform this biological function, is of great interest. The present paper addresses a question, how soft outer hair cells can amplify the vibration of the much stiffer basilar membrane. It shows that the elaborate structure of the cochlea, which supports multiple modes of motion, must be a key to the exquisite performance of the mammalian ear. It also shows that the properties of outer hair cells obtained from isolated cell preparations are compatible with their physiological function.

## Introduction

The mammalian hearing range may extend up to 100 kHz or beyond [1], depending on the species, quite remarkable for a biological system. Such function may call for a special mechanism aside from the unique “electromotile” machinery in outer hair cells (OHCs), which is essential for the performance of the mammalian ear [2].

Electromotility [3] is based on the piezoelectricity of the lateral wall [4]. As such, the response to direct mechanical stimulation to the cell body is intrinsically fast. However, this direct path is ineffective for the physiological function, and the effective response must depend on the mechanoreceptors in the hair bundle [5, 6]. Thus, the response must be mediated by the membrane potential, which is attenuated by the low-pass (*∼* kHz) nature of their intrinsic electric circuit, due to the capacitance of the cell membrane. In a system with resonance, this attenuation can be absent near the resonance frequency, where the membrane capacitance diminishes [7, 8].

The effectiveness of OHCs is also subjected to the impedance mismatch between soft OHCs and the stiff basilar membrane (BM), which is essential for frequency selectivity. If an OHC and the basilar membrane form a single oscillator, the upper limit of the effectiveness is *∼*10 kHz [5, 8], not high enough for covering the auditory range of mammalian hearing.

This limitation could be reduced if the OHCs and the BM are elements of separate oscillators and energy transfer between them is effective. Multiple modes of motion, which place OHCs and the BM in separate oscillators, are reported by recent experimental reports, using optical coherence tomography (OCT) [9–13]. In addition, there are theoretical models, which place OHCs between two inertial masses to explain the tuning curve of the ear [14–18]. Even though the neuronal output of the cochlea is better correlated with the movement of the reticular lamina [19, 20], BM motion is essential, being the basis for the tonotopic map of the cochlea [21].

All these studies, to my knowledge, do not address the effectiveness of energy transfer between oscillators at high frequencies, where impedance mismatch is the most pronounced [8]. The objective of the present study is to examine the effectiveness of energy transfer between the oscillators in the simplest cases of coupled oscillators. The models used retain the essential features of the cochlea together with the properties of OHCs determined from isolated cell preparations.

More specifically, we examine a set of two oscillators, heavy and light, coupled with either elastic or viscous element. First, a single mode oscillator with an OHC is described to illustrate the effect of impedance mismatch. Then systems of two coupled oscillators are examined. The light oscillator (LO) incorporates an OHC and the heavy one (HO) includes the BM. Coupling is either elastic or viscous. The OHC is stimulated by either the motion of HO or that of LO.

The present treatment focuses on local energy balance to address the upper bound of the auditory frequency. This condition is realized at locations, where energy influx and out-flux along the lateral direction are equal. It has been shown that such a condition is satisfied at any location, where the traveling wave of a given frequency stops [22].

### Single mode of motion

Let us start from a simple model oscillator system, into which an OHC is incorporated (Fig. 1). The equation of motion can be formally written as

**Figure 1:**
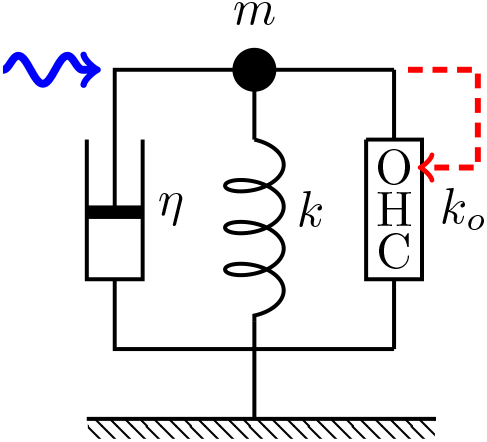
A single oscillator model. The oscillator consists of mass *m*, an external elastic element with stiffness *k*, a damper with drag coefficient *η*, and an outer hair cell (OHC), which responds to the movement of the mass (dashed red arrow). This system is driven by a sinusoidal waveform with angular frequency *ω* (wavy blue arrow).

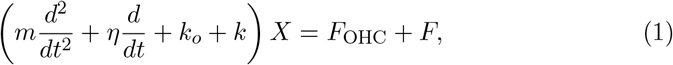

where *m* is the mass, *η* drag coefficient, *k* the stiffness of the external elastic load, *k*_*o*_ the material stiffness of the OHC, *X* is the length of the OHC, and *k*_*e*_ the stiffness of external load. *F* is an external force and *F*_OHC_ is the force generated by the OHC.

Consider the case, where a force *F* applied externally has a periodic waveform with angular frequency *ω*. We can write

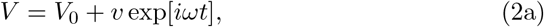

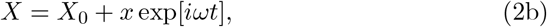

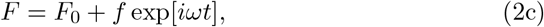

where *V* is the receptor potential, which is generated by the mechanosensitivity of the hair bundle.

In the frequency range, where capacitive conductance is greater than ionic conductance of the cell membrane and for a small amplitude *x*, the equation of motion can be expressed as [6]

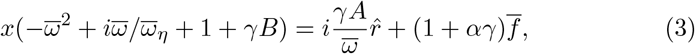

where 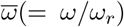 is the frequency normalized by the mechanical resonance frequency 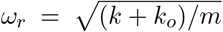 and 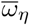 is normalized viscoelastic roll-off frequency. The quantity *γ* represents relative voltage sensitivity of prestin, the motile membrane protein of OHCs [23]. Its maximal value is set to unity (i.e. 0 ≤ *γ* ≤ 1) and it will be referred to as “the electromotile (em) parameter” or the “operating point parameter.”

The factor *A* is related to the force generation the response of the OHC through the mechanosensitivity of the hair bundle (HB) and *α* is a small factor (*<* 0.1) due to the direct piezoelectric mechanosensitivity of the cell body [6]. For this reason, the resonance in which OHC is involved is semi-piezoelectric rather than pure piezoelectric [6]. For this reason, this factor can be ignored. The factor *B* represents the stiffness change associated with conformational changes of the motile element. Their definitions are given in Appendix A. The quantity 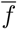 is given by

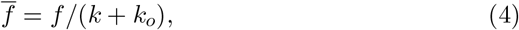

The quantify 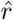 is the relative change of hair bundle resistance, which can be proportional to *x*. Thus, we may put 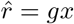. This substitution leads to

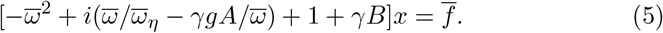

Notice that the factor *A* counteracts drag and factor *B* increases the stiffness of the cell due to strain-induced polarization [6]. Eq. 5 assumes that the mechanosensitivity of the piezoelectric cell body is smaller than that of the hair bundle, leading to minor underestimates of the amplitude (up to 10%)[6]. This assumption makes the connectivity, whether series or parallel, of the OHC unimportant and further facilitates the application of the present approach to coupled oscillator models.

The performance of the OHC in the system can be quantified by power gain *G*(*ω*), which is the ratio of power output to power input. Power output can be determined by the viscous dissipation *η* |*dX/dt*|^2^ because elastic energy is recovered during a full cycle. Power input can be expressed as Re[*F · dX/dt*].

Thus, power gain can be expressed by

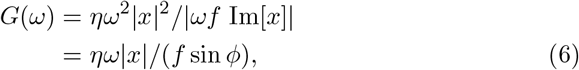

where *x* is given by Eq. 5, Re[…] and Im[…] represent, respectively, real part and imaginary part, and *ϕ* is the absolute value of the phase angle of *x* with respect to the external force. In the absence of the OHC, i.e. *A* = *B* = 0, we obtain *G*(*ω*) = 1 as expected.

## Coupled oscillator models

Consider a system with two harmonic oscillators, light and heavy. The light oscillator (LO) consists of an OHC and an elastic load *k*_*e*_ and inertia *m*. The displacement of this oscillator is *X*. The heavy oscillator (HO), which include the BM, consists inertia *M*, an elastic element with stiffness *K*, and viscous load with drag coefficient *η*. We assume, in general, that the two oscillators are coupled by an elastic element with stiffness *k*_*c*_ and a viscous element with drag coefficient *η*_*c*_. An external force *F* is applied to HO (Figs. 2 and 3). The motion of the two oscillators are described by

**Figure 2:**
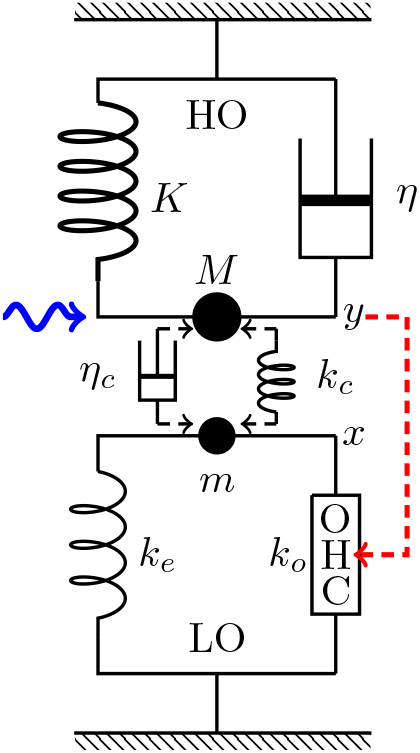
Coupled oscillators, in which OHC is driven by HO. HO (top) consists of mass *M*, an elastic element with stiffness *K*, and a damper with drag coefficient *η*. LO (bottom) consists of mass *m*, an elastic element with stiffness *k*_*e*_, and an OHC, which responds to the movement of HO (dashed red arrow). HO is driven by a sinusoidal waveform with angular frequency *ω* (wavy blue arrow) and the two oscillators are coupled either by an elastic element with stiffness *k*_*c*_ or by a damper with viscous coefficient *η*_*c*_.

**Figure 3:**
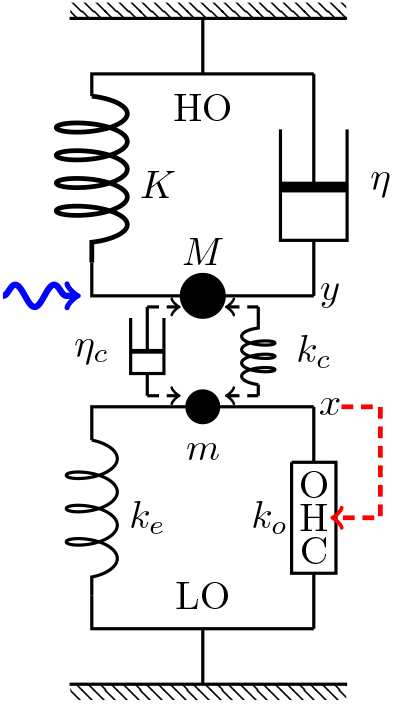
LO-driven coupled oscillators. HO consists of mass *M*, an elastic element with stiffness *K*, and a damper with drag coefficient *η*. LO (bottom) consists of mass *m*, an elastic element with stiffness *k*_*e*_, and OHC, which responds to the movement of LO (dashed red arrow). HO is driven by a sinusoidal waveform with angular frequency *ω* (wavy blue arrow) and the two oscillators are coupled either by an elastic element with stiffness *k*_*c*_ or a dashpot with viscous coefficient *η*_*c*_.

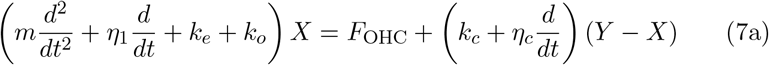

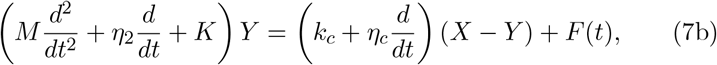

where *F*_OHC_ is active force exerted by the OHC, which is stimulated at its hair bundle. In addition, we assume that the main drag *η* belongs to HO. If we assume the opposite that the main drag belong to LO, the OHC cannot affect the oscillation of HO.

Classical analyses show that energy transfer between two coupled oscillators is rather complex despite the apparent simplicity of the equations [24, 25]. Here we examine only the response to continuous sinusoidal stimulation.

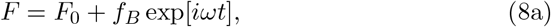

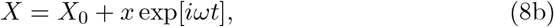

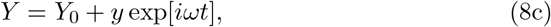

where *ω* is the angular frequency of stimulation. Notice also that the notation *f*_*B*_ is used for the amplitude of the external force in the systems of coupled oscillators. Now *F*_OHC_ can be expressed in a manner similar to in Eq. 5 using *A* and *B* because the force directly applied to the cell body is only of secondary importance for force production as discussed in the single mode oscillator.

In the case of viscosity coupling these quantities are defined by Eq. A1. In the case of elastic coupling, *A* and *B* need to be replaced respectively by *A*_*e*_ and *B*_*e*_, where the stiffness ratios are replaced due to the presence of an additional elastic element. Namely, *k*_*o*_*/*(*k*_*o*_ + *k*_*e*_) in *A* is replaced by *k*_*o*_*/*(*k*_*o*_ + *k*_*e*_ + *k*_*c*_) in *A*_*e*_ and *k*_*e*_*/*(*k*_*e*_ + *k*_*o*_) in *B* by (*k*_*e*_ + *k*_*c*_)*/*(*k*_*e*_ + *k*_*c*_ + *k*_*o*_) in *B*_*e*_.

In general, bending of the OHC hair bundle that results in 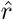 may depend on *x*, the displacement of LO, as well as on *y*, the displacement of HO. For simplicity, two extreme cases will be examined here: 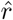 depends only on *y* (HO-driven mode), or only on *x* (LO-driven mode). Coupling may have both viscous and elastic components. However, we assume coupling is purely elastic or purely viscous for simplicity.

Power gain *G*(*ω*) is obtained analogous to Eq. 6 for the simple oscillator,

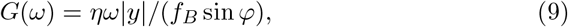

where *φ* is the absolute value of the phase angle of *y*. Notice that amplifier gain here is determined by the amplitude *y* of HO because both external force and drag work on HO.

### HO-driven case

We have assumed that the main drag of the system belongs to HO. If this drag is due to the shear in the subtectorial space, it may appear natural that HO motion stimulates the hair bundle of the OHC (Fig. 2), i.e. 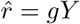.

#### Viscous coupling

Let *ω*_*η*_ be the viscoelastic roll-off frequency of HO. With new parameters defined by *s* = *K/k* and re-defined external force amplitude *f* = *f*_*B*_*/K*, the set of equations can be written as

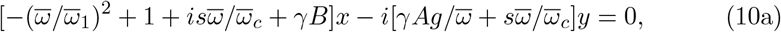

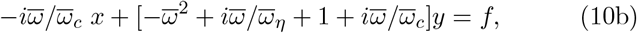

where *ω*_*r*_ is the resonance (angular) frequency of HO, i.e. 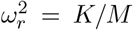, and 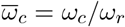 with *ω*_*c*_ = *K/η*_*c*_. All frequencies are normalized to *ω*_*r*_. For example, *ω* = *ω/ω*_*r*_. The quantity *ω*_1_ is the ratio of the resonance frequency of the LO to that of HO. Notice that *f* has the dimensionality of length.

### Elastic coupling

Coupling introduced by an elastic element *k*_*c*_ would elevate the resonance frequency of LO more than that of HO. The parameter *ω*_1_ adjusts the resonance frequency of LO.

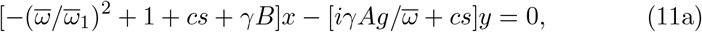

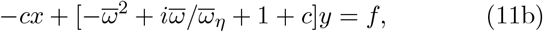

where *c* is defined by *c* = *k*_*c*_*/K*.

### LO-driven case

An alternative source of OHC stimulation is the displacement *x* of LO. This path of stimulation creates a direct feedback loop (Fig. 3).

#### Viscous coupling

The set of equations of motion can be written as

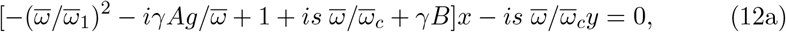

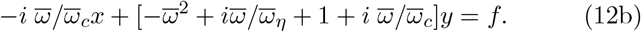

The phase relationship between *x* and *y* is determined by Eq. 12a.

#### Elastic coupling

The time dependent components follow the following equation

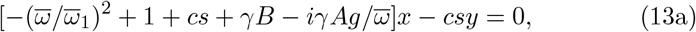

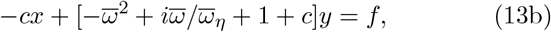

### Performance at high frequencies

To examine the performance of the coupled oscillator models at high frequencies, here we use the values of parameters that match the mechanical resonance frequency near 40 kHz (See Appendix B).

In comparing the performance of the single oscillator (SO) and coupled oscillators (COs), we use the following sets of parameter values:

For a single oscillator,

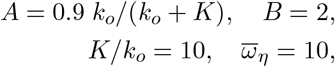

where *K* is the stiffness of the basilar membrane.

For coupled oscillators,

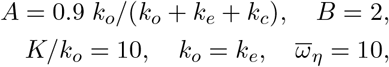

where *k*_*e*_ is the stiffness of the direct elastic load to OHC and *k*_*c*_ the stiffness of the elastic coupling element. For viscous coupling, *k*_*c*_ = 0. Adjustable parameters are 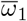 and a coupling parameter, either *c* = *k*_*c*_*/K* or 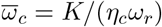 (See Figs. 2 and 3).

### Single mode oscillator

Let us start from the performance of the single mode oscillator before examining coupled oscillator models.

The steady state amplitude is obtained by solving Eq. 5. The amplitude is normalized by the input 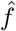 (Figs. 4). As *γ*, the operating point variable of the OHC, increases, the peak amplitude shifts to higher frequencies independent of the elastic load (Fig. 5B). The peak height, however, does not increases with *γ* with elastic load of *K/k*_*o*_ = 10. Only if the stiffness ratio is reduced to 2.4 or lower, the peak height does increase Fig. 5A).

**Figure 4:**
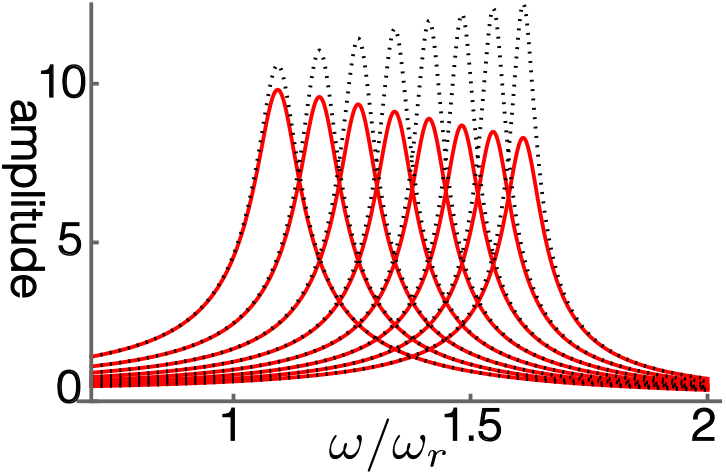
Amplitude of single mode oscillator plotted against frequency (normalized to *ω*_*r*_). Peak values are replotted in Fig. 5A. Stiffness ratio of load to OHC is 10, assuming the BM is the elastic load (red). The result of elastic ratio of 5 (dotted) is shown for comparison. *γ* (operating point variable) runs from 0.03 (left) to 0.24 with increment of 0.03. The unit of the ordinate axis is *f* (= *F/K*).

**Figure 5:**
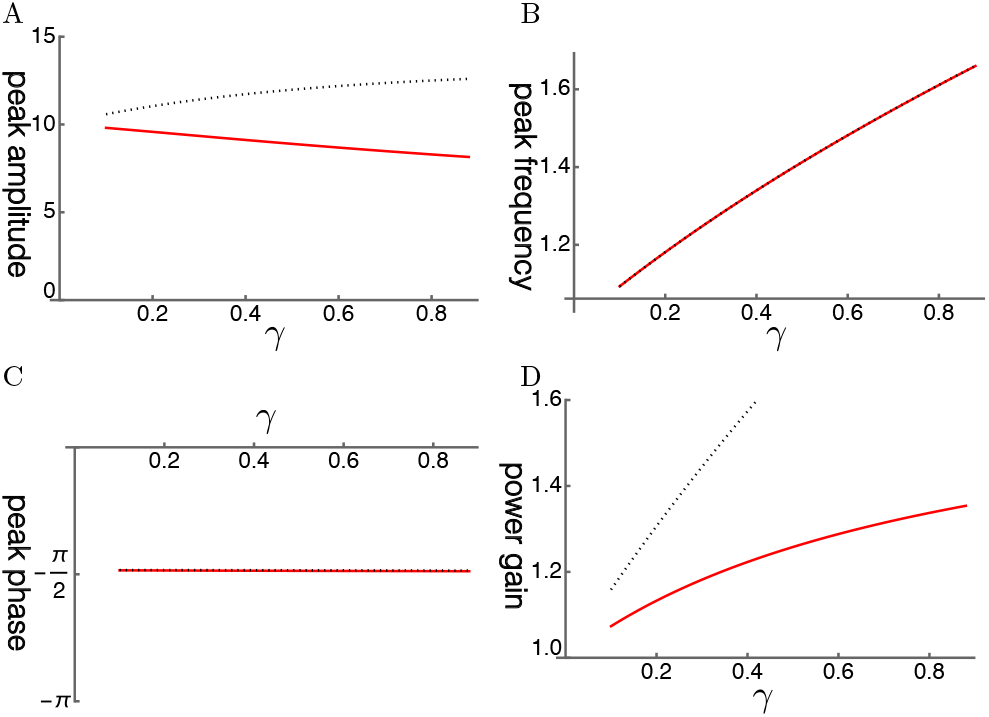
A: Peak amplitude, B: Peak frequency (normalized to the resonance frequency), C: peak phase, and D: Power gain (ratio of power output to power input) are respectively plotted against *γ*, the operating point variable. The stiffness ratio is 10 (red), reflecting the value of *K/k*_*o*_. Stiffness ratio of 5 (dotted) is shown for comparison. The peak phase is close to −*π/*2, consistent with resonance.

The phase of the oscillator lags by *π/*2 as expected from the maximal amplitude of the damped oscillator (Fig. 5C). That makes power gain similar to the amplitude of oscillation (Fig. 5D). The low power gain is significantly attenuated by input impedance mismatch.

### Coupled oscillators

In the following, we will examine if OHCs can be more effective in coupled oscillators than in the single mode oscillator. The coupled oscillator models have more parameters, which include stiffness ratio *s* and resonance frequency ratio *ω*_1_ of the two oscillators, and coupling parameters (elastic element *c* and viscous element *η*_*c*_).

Conditions for large gain in amplitude are sought in all four cases. It turned out that large gains can be obtained in the systems with elastic coupling. In the following, examples are shown of the systems with elastic coupling. These examples are typical ones but they do not necessarily show all shared features of the conditions.

Viscous coupling is also capable of producing amplification even though their gain is smaller elastic coupled coupling. Examples of cases with viscous coupling are shown in Supplementary Material.

### Elastically coupled HO-driven cases

The HO-driven mode is interesting because we assume that HO is associated with the dominant drag, which likely stems from the shear in the subtectorial space and this shear should stimulate the hair bundle of the OHC (Fig. 2).

With elastic coupling, it is possible to find conditions, under which the amplitude gain of the LO reaches *∼*20 fold over the single mode oscillator (Fig. 6A). However, the amplitude gains reach their maximal values before *γ*, the operating point parameter, reaches unity. The peak frequency is about the same for the two oscillators and increases with the operating point variable (Fig. 6B).

**Figure 6:**
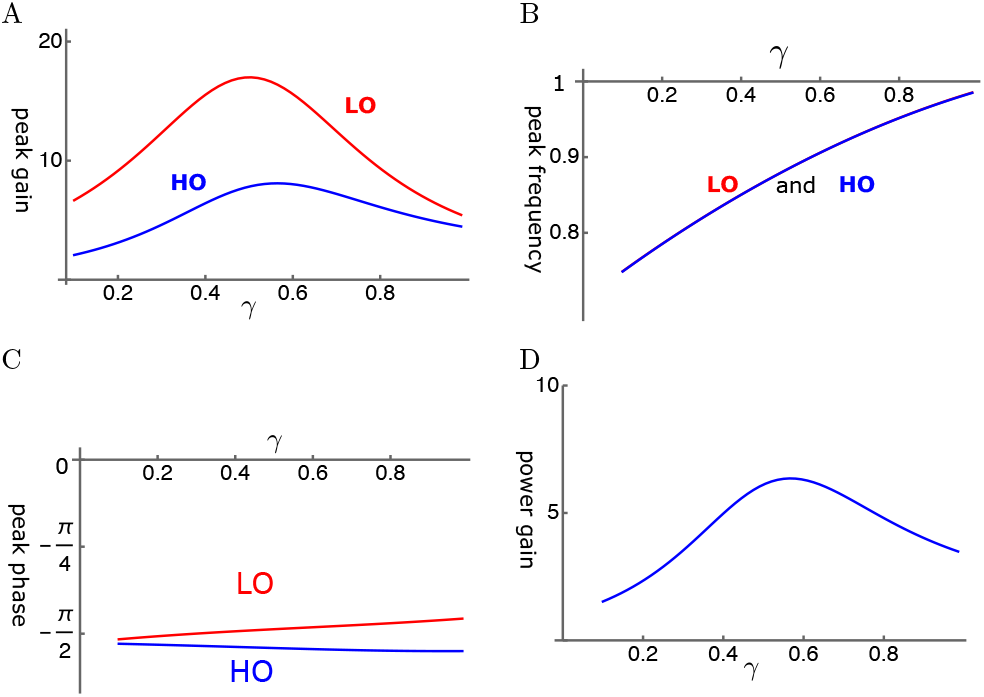
HO-driven elastically coupled oscillators (HOE). A: Amplitude gain over SO mode, B: Peak frequency, and C: The phase with respect to the external force, of each oscillator are plotted against *γ*. LO (red) and HO (blue). D: Power gain. The set of parameter values: *c*=0.2, *ω*_1_=0.29 in Eq. 11.

The phase of LO is ahead of HO, consistent with the amplifying role of LO. However, the phase difference is rather small (Fig. 6C). The power gain coincides with the amplitude maximum of HO and reaches about 6 at the peak (Fig. 6D).

### Elastically coupled LO-driven cases

Here we assume that the OHC is primarily driven by the motion of LO rather than that of HO (Fig. 3). With elastic coupling, the amplitude gain can exceed 100 fold for LO and 80 fold for HO (Fig. 7A). The amplitude gain generally increases with the operating point variable *γ* as expected. This increase can be monotonic as in Fig. 7A. However, it can peak before *γ* reaches unity, similar to HO-driven cases, depending on the set of parameter values.

**Figure 7:**
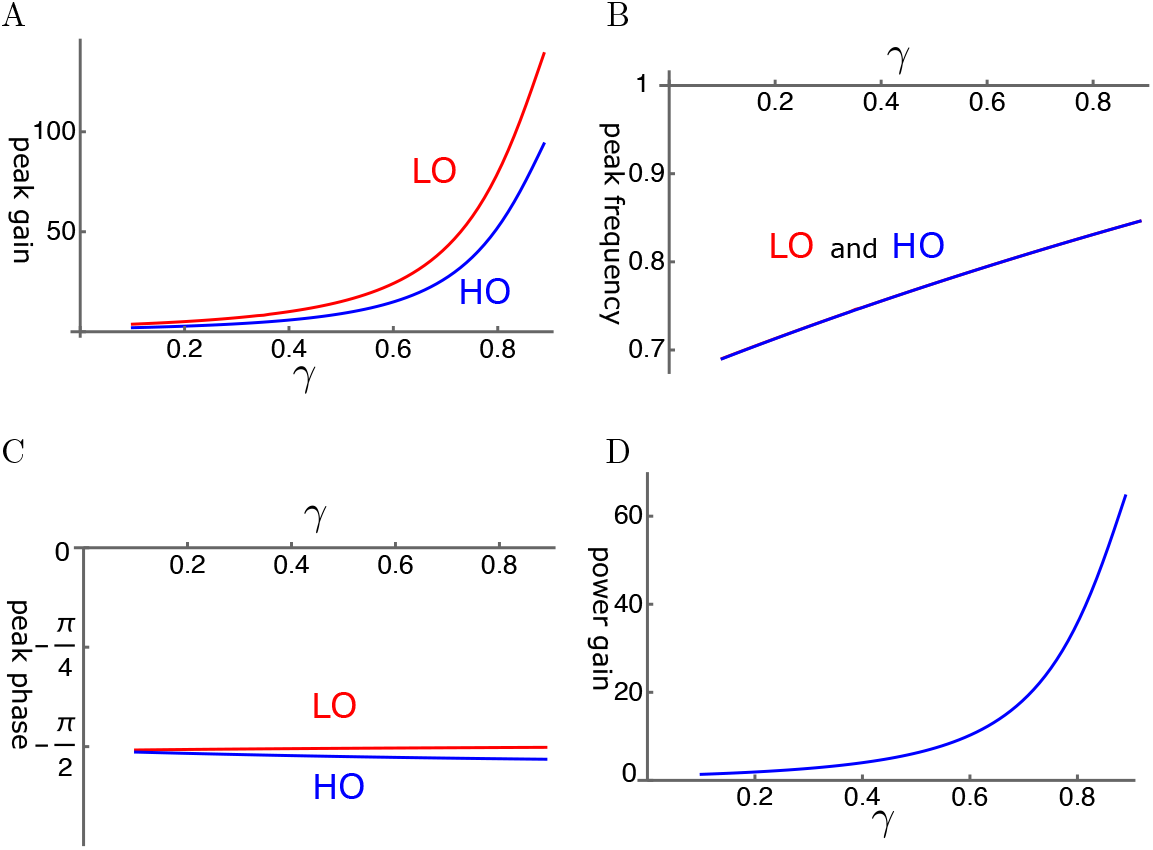
An example of LO-driven elastically coupled oscillators (LOE). A: Amplitude gain over SO mode, B: Peak frequency, and C: The phase with respect to the external force, of each oscillator, is plotted against *γ*. LO (red) and HO (blue). D: Power gain. The set of parameter values: *c*=0.6, *ω*_1_=0.18 in Eq. 13.

The peak frequencies are similar for both oscillators (Fig. 7B). The LO is ahead of HO in phase, indicating the amplifying role of the LO even though the difference is rather small (Fig. 7C). The effect of the LO in affecting the amplitude of LO is quite large despite its much smaller mass and stiffness. The power gain reaches about 60 where the amplitude gain of HO is maximal (Fig. 7D).

The frequency dependence shows quite sharp tuning together with the large amplitude gain (Fig. 8).

**Figure 8:**
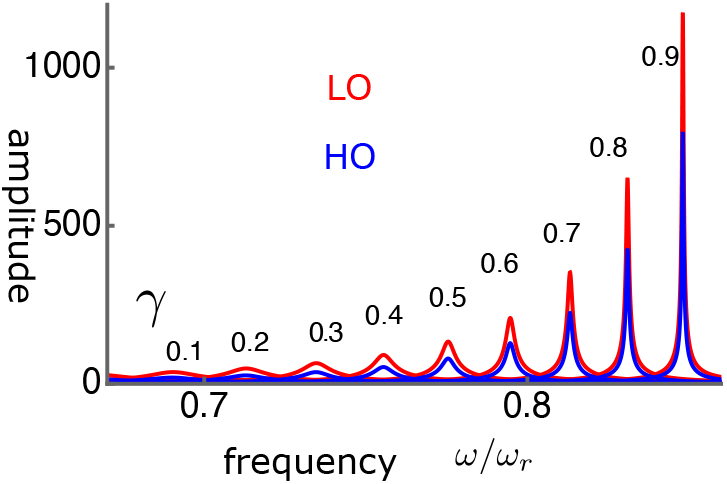
The amplitudes LO (red) and HO (blue) in LO-driven elastically coupled (LOE) oscillators. The peak values are plotted in Fig. 7A. The abscissa is the frequency normalized to *ω/ω*_*r*_. The unit of the amplitude (ordinate axis) is *f/K*. The values of *γ* are from 0.1(left) to 0.9(right) with an increment of 0.1.

### Interference of coupling

The case of LO-stimulation with elastic coupling (LOE mode) is the most effective in utilizing OHC. It leads to about 60 fold power gain (Fig. 7D) and amplitude gain of about 80 fold for HO (Fig. 7D). The case of HO-stimulation with elastic coupling (HOE mode) may lead to a relatively good amplitude gain for LO, even though it is less effective than LOE. In view of similar phase relationships of the oscillators in those two cases, a superposition of these two stimulation modes could be constructive.

## Discussion

The numerical examination presented here concerns only high frequency performance. At lower frequencies, such as 10 kHz, viscosity coupled cases could produce reasonable gain. However, a high amplifier gain may not be a critical condition at such frequency ranges.

### Input impedance mismatch

The coupled oscillator models can make OHCs more effective by reducing the constraint of impedance mismatch. However, the performance of coupled oscillator models significantly varies depending on the type of coupling and how OHCs are stimulated.

More specifically, optimal amplification of this system by the OHC is achieved in the case where coupling is elastic and the OHC is stimulated primarily by the oscillation of LO. Under such conditions, the gain is not impeded by a large stiffness mismatch between the OHC and the BM.

### Some characteristics

Detailed properties of coupled oscillators vary depending on the set of parameter values. For large amplifier gain, the following observations can be made.

### Frequency dependence

Strong coupling is required for making OHCs effective. In the case of elastic coupling, for example, the stiffness of the coupling element must be similar to that of the elastic element of LO. This condition appears to lead to another feature that the resonance frequency of LO is somewhat lower than that of HO. With increased OHC electromotility (increased *γ*), the resonance frequency tends to increase, reminiscent of the half-octave shift [26, 27].

### Stability and *γ*-dependence

Amplitude gain of a coupled oscillator is in general an increasing function of *γ*, the operating point as expected. Some of the cases (See Fig. 7A) show monotonic increase with *γ* similar to the single mode oscillator with low elastic load (See Fig. 5A). However, depending on the condition, it can peak before the operating point variable *γ* reaches unity (See Figs. 6A). Such peaking disappears by reducing the value of *B*, which shifts the resonance frequency.

We could speculate that such a behavior could explain a report that lowering prestin density in OHCs by 34% does not have any reduction in the sensitivity of the ear [28]. That is because both *A* and *B* are proportional to *N*, the number of motile units in an OHC (See Table A.1). Thus, *N* and *γ* have the same effect on these parameters.

Amplitude gains are very sensitive to small variations of the cochlear parameters. Such a sensitivity may result in uneven power gain. Amplitudes can be smooth functions of the operating point variable *γ* as shown in the figures. They can also show singular dependence on *γ*, indicating spontaneous oscillation. These observations could have some bearing on otoacoustic emissions [29].

### Structural implications

The behavior of the equations examined must have a structural basis, including the nature of coupling elements, the effect of cell length, and movements in the subtectorial space.

### Nature of coupling

The cell body of OHCs is held by the reticular lamina at the apical end, forming tight junctions, and by Deiters’ cup at the basal end. This structure can introduce viscoelastic interaction with Deiters’ cells [30], which serves as a basis for viscous coupling [17, 31]. The present analysis does not show that viscous coupling is effective to enhance the amplitude of LO at high frequencies. Presumably OHCs are held elastically, at least in the basal high-frequency region of the cochlea.

### Cell length

Short hair cells are more effective, having larger values of amplifying parameter *u* because of their smaller values of *C*_0_ and larger values of *k*_*o*_. Indeed, basal cells, which operate high frequencies, are short. For example, basal cells in both guinea pigs and mice are short (≲ 20 *µ*m long), even though their length gradient is steeper for guinea pigs [32].

### Subtectorial drag and OHC stimulation

The models examined in the present treatment assume that the main drag of the system is associated with HO. Such a condition could be realized if the main drag imposed on HO is outside the subtectorial gap.

However, the most likely source of the main drag of the system could be the shear in the subtectorial space between the tectorial membrane and the reticular lamina [33]. This picture indicates that the shear is associated with the motion of HO. If we accept that hair bundle bending is associated with this shear, the OHC should be stimulated by the displacement of HO. That leads to HO-stimulation models.

Can the hair bundle of the OHC be primarily stimulated by LO without incurring significant drag? Such a condition could be realized if LO moves the hair bundle in the direction perpendicular to the reticular lamina, resulting in effective bending of the hair bundle. However, the movement of LO should not incur significant drag. It would be possible that displaced water volume could be accommodated by local displacement of the TM owing to its pliability [34] and mechanical anisotropy [35, 36]. Then, the movement of LO does not result in viscous drag because it does not lead to fluid flow along the gap.

### Speed of the motile element

We assumed that prestin, the motile protein that drives OHCs, undergoes conformational transitions fast enough so that mechanical constraints determine the frequency dependence. This assumption is consistent with the experimental data on isometric force generation by OHC [37] and current noise spectrum of OHC membrane [38]. It is also in line with a recent analysis that movement of organ of Corti measured with OCT is consistent with the cycle-by-cycle force application [39].

However, the frequency of conformational changes must have an upper bound. Recent repots that the roll-off frequencies of the voltage-dependent component of OHC membrane capacitance suggest 30 kHz, [40] higher than older values of up to 20 kHz [41, 42]. Those gating frequencies reported could reflect extracellular factors, such as viscoelastic process, of their experimental configuration.

The present study provides a new perspective: with a large gain obtained by coupled oscillators, this issue is not so important. With a finite gating frequency *ω*_*g*_, the amplitude is attenuated by a factor 1*/*(1 + *ω*_*g*_*/ω*) [43]. If the gating frequency is 20 kHz, this attenuation factor is 1/3 at 40 kHz, a small fraction of the expected gain.

### Implications to cochlear models

Cochlear models that are proposed to explain the performance of the mammalian ear, particularly the tuning curve, can be classified into two groups: microscopic and macroscopic models.

Numerous recent theoretical models do assume multiple degrees of freedom [44]. These models are of particular interest in recent years because they can be related to recent OCT experiments. These models can, in principle, overcome the issue of input impedance by choosing suitable parameter values because of their models have multiple degrees of freedom.

Macroscopic models of cochlear mechanics [45] also are of interest because they assume that OHCs apply force directly on the BM. Such models ignore issues resulting from microscopic structure, such as low-pass characteristics of intrinsic cellular electric circuit and impedance mismatch.

Interestingly, however, the present analysis of coupled oscillator models shows a resemblance to the assumptions of those macroscopic models: The amplifier gain is not seriously constrained by the impedance mismatch. The difference in the amplitude and in the phase of the two oscillators is rather small. This observation does show the viability of these macroscopic models as giving approximate descriptions. However, it may not indicate the physical details in the inner the ear.

## Conclusions

A large impedance mismatch between the BM and the OHC impedes energy transmission from the softer OHC to the oscillation of the stiffer BM if the BM and the OHC are in the same oscillator. However, if these elements are incorporated into separate oscillators that are coupled, a significant improvement in the efficiency of energy transmission can be achieved.

Among the modes of motion examined, the system of elastically coupled with LO-stimulated OHC is most effective in utilizing the OHC as the amplifier. Under optimal conditions, both amplitudes and phases of these oscillators are close because these oscillators are strongly coupled.

The models examined here are the simplest possible cases. The real ear would be much more complex and can have more modes of motion. Nonetheless, these simple model systems could provide some insight into the working of the real system. It is likely that multiple modes of motion supported by the complexity of the organ of Corti are essential for the performance of the mammalian ear by making OHCs effective as the amplifier.

## Acknowledgments

I thank Drs. Richard Chadwick and Catherine Weisz of NIDCD for helpful comments. Dr. Chadwick pointed out the significance of Eq. 28 of Wang et al [22]. I also appreciated help by Drs. Matthew Kelley and Inna Belyantseva for reading the manuscript. This research was supported in part by the Intramural Research Program of the NIH, NIDCD.

## Appendix A Definitions of Parameters

For the system that corresponds to Fig. 1, the factors *A, B*, and *α* are given by [6]

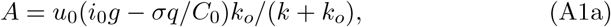

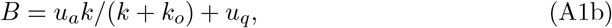

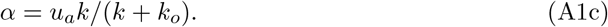

Quantities *u, u*_*a*_, and *u*_*q*_ represent hair bundle sensitivity and piezoelectricity of the OHC. More specifically, *u* represents mechanoelectric coupling, *u*_*a*_ mechanical term, and *u*_*q*_ electrical term, while all of them depend on hair bundle sensitivity. See Table A.1 for definitions.

The elastic load *k* depends on the configuration. For single mode oscillator (SO), *k* = *K*, for coupled oscillators, *k* = *k*_*e*_ + *k*_*c*_, where for viscous coupling, *k*_*c*_ = 0 (See Figs. 2 and 3).

**Table A1:** Parameter definitions.

| symbol | definition | parameter | definition |
| --- | --- | --- | --- |
| $P_0$ | motor conformation* | $\beta$ | $1/(k_B T)$ |
| $a$ | prestin displacement | $\gamma$ | $4P_0(1 - P_0)$ |
| $\omega_r$ | resonance frequency | $u_a$ | $\beta a^2 k_o N/4$ |
| $q$ | prestin charge | $u_q$ | $\beta q^2 N/(4C_0)$ |
| $N$ | number of prestin | $u_0$ | $\beta q/(4\omega_r C_0)$ |
| $C_0$ | cell capacitance | $\omega_\eta$ | $K/\eta$ |
| $i_0$ | resting current | $\omega_1$ | resonance frequency ratio |
| $k_o$ | cell stiffness | $\hat{r}$ | relative change of HB resistance |
| $k_e$ | parallel elastic load | $\sigma$ | membrane conductance |
| $k_c$ | coupling spring | $f_B$ | external force (amplitude) |

Note: $k_B$ is Boltzmann's constant, $T$ the temperature. \* The motile activity of OHCs is described by a two state model. $P_0$ is the fraction of one of the two states in a single cell at the operating point. In the absence of elastic coupling $k_c = 0$

## Appendix B Parameter values

To find a set of adequate parameter values for a high frequency (40 kHz) location of guinea pigs, the parameter values are examined for a more readily obtainable 20 kHz location, and then they are extrapolated to 40 kHz location.

### Cellular factors (20 kHz)

For a 20 *µ*m-long cell, typical of the 10 to 20 kHz cochlear region, we have for the linear capacitance *C*_0_ = 8 pF and the maximal electromotile displacement *aN* = 1*µ*m, which is 5% of the resting cell length. Most in vitro experiments show the unitary motile charge of *q* = 0.8*e*, where *e* is the electronic charge. The resting membrane potential is near the optimal range for the motile element. The resting basolateral resistance is 7 MΩ and the resting membrane potential of −50 mV requires the resting apical resistance of 30 MΩ. These values lead to *i*_0_ = 4 nA. These parameter values are summarized in Table B.1

The stiffness *k*_*o*_ of a 20 *µ*m-long OHC is about 20 mN/m, given the specific stiffness of 510 nN/m per unit strain [46]. However, this value may have some uncertainty. The bottom part, up to 10 *µ*m, of the OHC is held by the Deiters’ cup. If this structure works as a damper [31], the length of elastic displacement is larger because the displacement includes the part within the cup. If, on the contrary, the structure is tight and rigid, not allowing slippage, the value of *k*_*o*_ must be higher.

The parameter values in Table B.1 lead to a set of values for an OHC at 20kHz location:

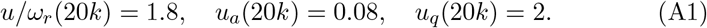

The second term containing *σ* in the coefficient *A* is not significant at 20 kHz location because the ratio *i*_*o*_*/σ* is small unlike at 4 kHz location [6]. That is because HB conductance, which primarily determines *i*_0_, increases much sharper than the cellular conductance toward the base. The coefficient *B* is approximately 2. It is not sensitive to the ratio of the load ratio because it *u*_*a*_ is smaller than *u*_*q*_.

**Table B1:**
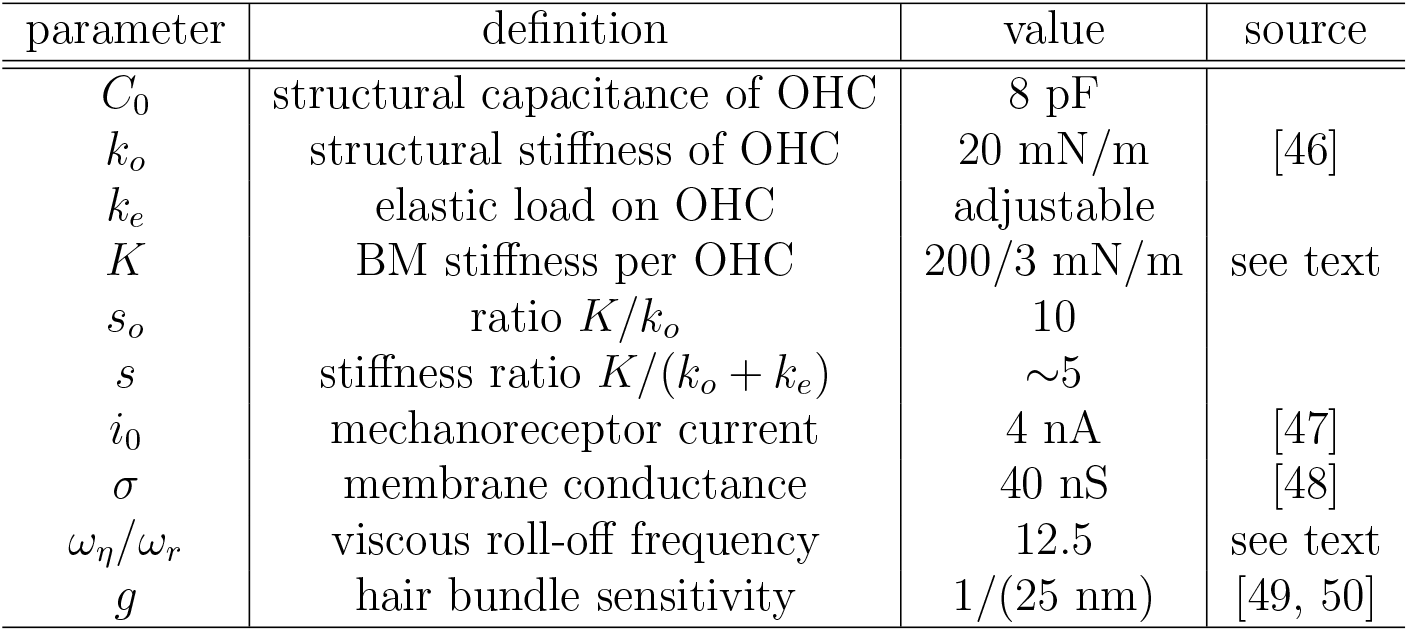
Parameter values at 20 kHz location.

### Cochlear factors (20 kHz)

Now assume that the BM is the source of the stiffness *K*. The best frequency location for 20 kHz in guinea pigs is 3 mm from the stapes according to the Greenwood function, assuming the length of the BM is 18.5 mm [51, 52]. The stiffness of the BM at this location is about 0.21 N/m measured with a probe with 25 *µ*m diameter tip [53]. This value is compatible with the ones obtained with a probe with 10 *µ*m tip [54]. Thus, the stiffness ratio *s*_*o*_ is *∼*10.

The intrinsic mechanical resonance frequency of the location is somewhat uncertain because of the so-called “half-octave shift” [55]. If the intrinsic mechanical resonance corresponds to the “passive” condit<u>i</u>on, the resonance frequency of the 20 kHz location is 14 kHz 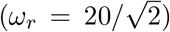 being a half octave lower. However, it could be the opposite because viscous damping brings the peak to a lower frequency if it is not counteracted.

A significant contribution to friction is expected from the gap between the tectorial membrane and the reticular lamina. The friction coefficient of this gap can be estimated by a formula *µS*^2^*/d*, where *µ* is the viscosity of the fluid, *S* the area of the gap per OHC, and *d* the gap, provided that the thickness of the boundary layer, which is *∼* 3.6 *µ*m for 20 kHz [56], is greater than the gap. If we assume *S* is 10*µm×* 15*µm* and *d* 1*µm*, the friction coefficient is 1.2 *×* 10^−7^ N/m [5, 8]. With the resonance frequency of 14 kHz, the gap friction leads to a value 12.5 for *ω*_*η*_*/ω*_*r*_.

### Extrapolation to higher frequencies

Of the OHC parameters, *B* does not depend heavily on the resonance frequency because it is dominated by the electrical term *u*_*q*_ (See Eq. A1b). The amplification parameter *A* decreases at higher frequency locations because it is inversely proportional to the resonance frequency *ω*_*r*_. However, such a reduction can be compensated by other factors: OHCs at higher frequency locations have larger resting current *i*_0_ owing to larger hair bundle conductance and larger structural stiffness *k*_*o*_ of the cell body owing to its shorter cell length (See Eq. A1a and Table A.1).

At higher frequency locations, the stiffness ratio *s* is expected to increase because the stiffness of the BM would increase more than the stiffness of OHCs. However, we proceed by assuming that the ratio *s* remains the same. The reason is an uncertainty in the effective length of OHCs. The stiffness is inversely proportional to the effective length of the lateral membrane, which is harder to determine with shorter OHCs because the connectivity in Deiters’ cup is ambiguous as described earlier. In addition, somewhat higher values for *s* do not lead to significantly different results.

To examine high frequency performance aiming at 40kHz, twice higher than 20kHz, where parameter values are examined above, a numerical analysis is performed, assuming the following set of parameter vales:

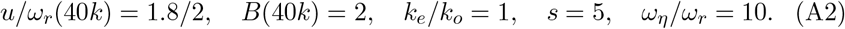

## Supplementary Material

Here viscosity coupled oscillators are presented. Each example is obtained after attempts of optimizing the amplitude gain. These cases do show amplification, but not as large as elasticity coupled cases.

### Viscosity coupled HO-driven case

The Deiters’ cup that links an OHC with the BM via Deiters’ cell could provide viscous coupling due to its morphology. For this reason, it is of interest to examine the effect of viscous coupling.

Even though viscous coupling of the oscillators can have an amplifying effect, it does not appear to be effective. Amplitude gain tends to be small (Fig. S1A). Peak frequency of LO and that of HO are close to each other except for at small values of *γ* (Fig. S1B). The phase of HO is ahead of LO (Fig. S1C). Since the force LO applies to HO is by *π/*2 ahead of the phase of LO, LO amplifies the motion of HO. Power gain is rather small (Fig. S1D) as expected from the amplitude of HO.

**Figure S1:**
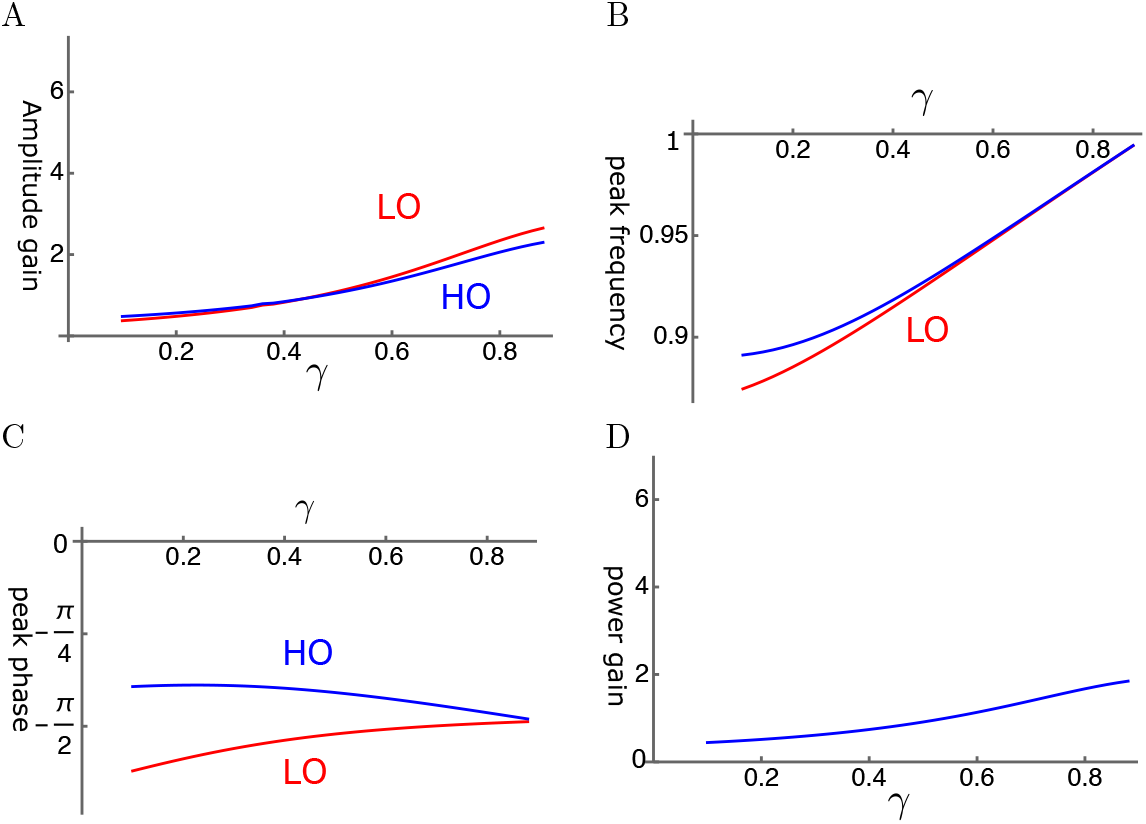
HO-driven viscosity coupled oscillators (HOV). A: Amplitude gain over SO mode, B: Peak frequency, and C: Peak phase with respect to the external force of each oscillator are plotted against *γ*, the operating point variable. LO (red) and HO (blue). D: Power gain is closely associated to HO amplitude. The set of parameter values: 1*/ω*_*c*_=0.3, *ω*_1_=0.33 in Eq. 10.

### Viscosity coupled LO-driven case

The amplitude gain for both LO and HO increases with the operating point variable *γ* (Fig. S2A). HO has somewhat higher peak frequency than LO for small *γ* (Fig. S2B). The phases of HO and LO are both close to *π/*4 and HO tends to be ahead of LO except for where *γ* is small (Fig. S2C). The power gain is not very large.

**Figure S2:**
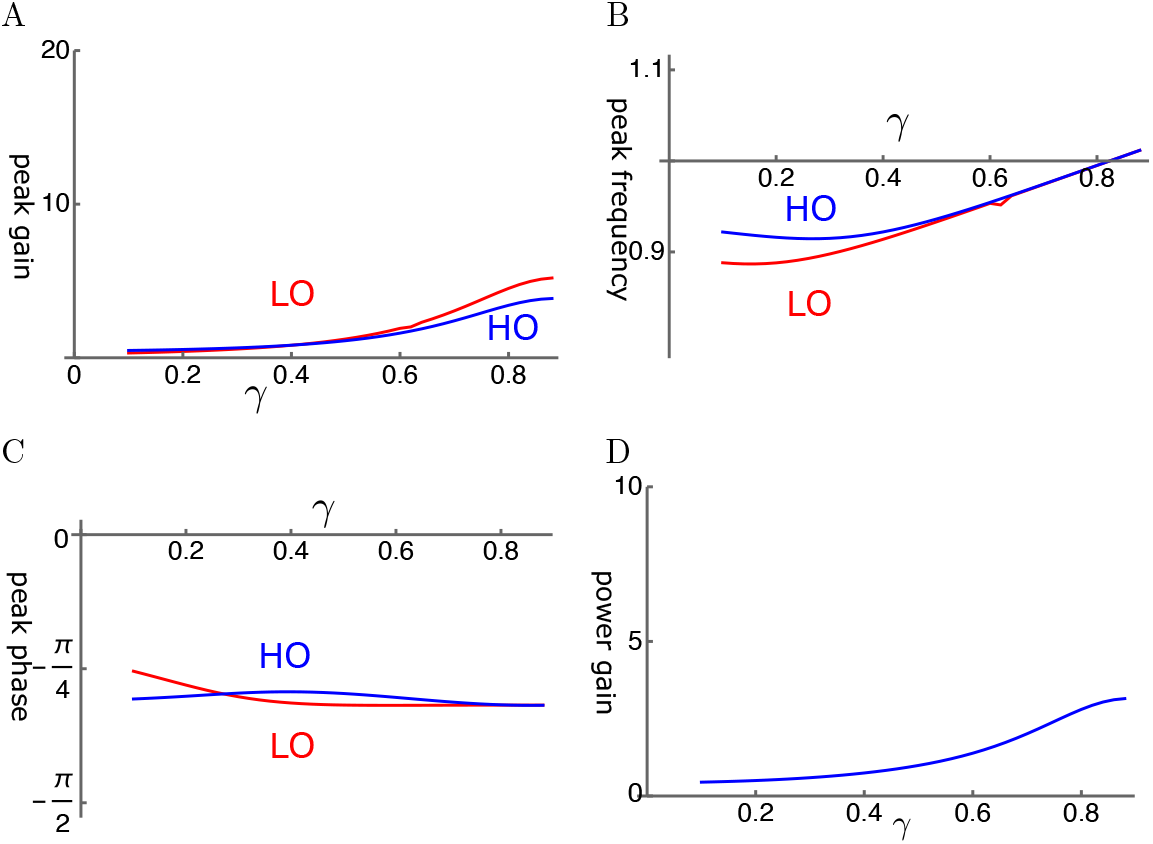
LO-driven viscosity-coupled oscillator (LOV). A: Amplitude gain over SO mode, B: Peak frequency, and C: The phase with respect to the external force, of each oscillator are plotted against *γ*. LO (red) and HO (blue). *η*^*c*^ = *η*_*c*_*ω*_*r*_*/K*. D: Power gain. The set of parameter values: 1*/ω*_*c*_=0.2, *ω*_1_=0.36 in Eq. 12.

